# Fish are poor sentinels for surveillance of riverine AMR

**DOI:** 10.1101/2024.10.22.619632

**Authors:** Faina Tskhay, Christoph Köbsch, Alan X. Elena, Johan Bengtsson-Palme, Thomas U. Berendonk, Uli Klümper

## Abstract

Effective surveillance of antimicrobial resistance (AMR) in the environment is crucial for assessing the human and animal health risk of AMR pollution. Wastewater treatment plants (WWTPs) are one of the main sources of AMR pollutants discharged into water bodies. One important factor for assessing the risks associated with such pollution is the colonization potential of the resistant bacteria (ARB) and resistance genes (ARGs) from the environment into human or animal microbiomes upon exposure. This study explores whether fish can act as sentinels for surveillance of AMR pollution in general and specifically the human colonization potential of ARB in rivers impacted by WWTP effluents. Two riverine fish species, Brown trout, and European bullhead, were sampled up- and downstream a German WWTP. The two fish species were chosen due to their different lifestyles: Trout are mainly actively swimming in the water phase, while bullheads are sedentary and river sediment-associated. The bacterial microbiomes and resistomes of fish gills, skin, and feces were compared with those of the respective river water and sediment up- and downstream of the WWTP. Microbiomes of both fish mirrored the changes in river water and sediment downstream of the WWTP, with significant shifts in bacterial community composition, particularly an increase in Proteobacteria and Verrucomicrobia. However, increases in ARG abundances observed in water and sediment downstream of the WWTP were not reflected in any of the fish-associated resistomes. This indicates that while the fish microbiome is sensitive to environmental changes, resistomes of poikilothermic animals such as fish are less responsive to colonization by ARB originating from WWTPs and may not serve as effective sentinels for assessing AMR pollution and colonization risks in freshwater environments. This study highlights the complexity of using wildlife as indicators for environmental AMR pollution and suggests that other species are better suited for surveillance efforts.

**Highlights:**

- WWTP effluent affected downstream river microbiome composition of water and sediment
- WWTP effluent significantly increased the level of six tested ARGs downstream
- Changes in environmental microbiome composition were mirrored in trout and bullhead
- No effect of the WWTP effluent on resistomes and individual ARG abundance in fishes

## Introduction

Excessive and improper use of antibiotics has facilitated rapid development of a global antimicrobial resistance (AMR) crisis. With almost five million deaths annually associated with resistant bacteria, AMR has become one of the biggest threats to human health [1]. Combating this global AMR crisis requires a comprehensive strategy integrating human health, animal health, and the environment. This strategy, known as the “One Health” concept, proposes surveillance as an essential component for mitigating the AMR problem [1,2]. However, despite the environment’s crucial contribution to the emergence and dissemination of AMR [3–6], current surveillance strategies focus primarily on clinical, food, and agricultural settings [7,8]. Meanwhile, wastewater treatment plants (WWTPs) are commonly described as one of the primary sources of anthropogenic antibiotic and AMR pollution to the environment [4,9–11]. Surveillance of such environmental hotspots can help estimate both the community AMR carriage and the impact of anthropogenic pollution at the direct intersection between the human and environmental spheres [12,13].

While modern wastewater treatment plants have been designed to remove solids, nutrients (e.g., nitrogen and phosphorous), and toxic and volatile compounds [14], their efficiency in eliminating pharmaceuticals including antibiotics still needs improvement [4,10,15]. In addition, WWTPs are not optimized to completely remove bacteria (including resistant ones) and antibiotic resistance genes (ARGs) [10,11,16]. Consequently, WWTP effluents contain significant amounts of antimicrobial-resistant bacteria (ARBs) and ARGs that are released into water bodies [4,17]. For example, WWTPs in Germany discharge an average of 3.30 × 10^14^ ARB cell equivalents daily [18]. This substantial load regularly increases the absolute and relative levels of detectable ARGs and ARBs in receiving waters [19–21] and sediments downstream of WWTPs [9,22]. However, if such increased AMR loads are connected to an increased risk of colonization of human and animal microbiomes exposed to the waters needs to be evaluated, since this is a crucial factor in the risk assessment during AMR surveillance [7]. Especially, whether this impact on the environmental microbial communities is also reflected in the microbiomes of riverine wildlife living downstream of WWTPs, remains largely unknown.

Considering the integral role of fish in various ecological niches, and their ubiquity in the aquatic environment, this study aims to determine if the fish microbiome can serve as a sentinel for the colonization potential of AMR introduced into water bodies impacted by WWTP effluent for future risk assessment during surveillance. Hence, in this study, we explored if changes in the environmental resistome caused by WWTP effluent could also be observed in fish. Fish interconnect all three ecological compartments of the “One Health” concept [23]. In the environment, fish are one of the most available and abundant living substrates in water that can capture and be colonized by ARB and ARGs. Due to their mobility, fish can act as carriers of resistance and transport ARB and ARGs over long distances [24]. Ultimately, ARB and ARG potentially carried by fish can be transferred to humans through fishing and subsequent consumption of the caught fish. To test whether the fish resistome does indeed reflect WWTP- induced changes to the rivers’ water and sediment resistomes, we compared the microbiome and resistome composition of gills, mucosal skin, and feces of two freshwater fish - Brown trout (*Salmo trutta*) and European bullhead (*Cottus gobio*) - obtained in a German rural river, either up- or downstream of the lone WWTP in that catchment. To further understand if such potential effects on the fish microbiome or resistome are general to all fish or depend on their lifestyle, we chose two fish species that differ by lifestyle. Trout actively swim in the river water column and can travel long distances, especially during spawning seasons [25,26]. In contrast, bullheads are predominantly sedentary, sediment-associated fish, exhibiting low movement behavior [27,28].

## Material & Methodology

### Sample collection

River water, sediment, and fish samples were collected in May 2022 from the Rauner Bach-Weiße Elster river catchment in the Vogtland region of Saxony, Germany. Additionally, wastewater effluent samples were collected from the municipal wastewater treatment plant (WWTP) discharge pipe into the river in the town of Adorf. The Rauner Bach, a right-bank tributary of the Weiße Elster, is located approximately 7.6 kilometers upstream of the Adorf WWTP and was selected as a site unaffected by wastewater effluent (50°16’33.1"N, 12°17’41.5"E). This stream, part of the “Rauner-und Haarbachtal” nature reserve, is characterized by low anthropogenic influence and a favorable conservation status for the bullhead (*Cottus gobio*) as an animal species of community interest according to the Habitats Directive [29]. The stream thereafter becomes the Weiße Elster River without any physical barriers. The WWTP discharges treated wastewater into the Weiße Elster River 3.6 kilometers before the downstream sampling site (50°20’38.9"N, 12°14’33.5"E). The downstream sampling site is notable for its significant bullhead population and is a popular spot for recreational fishing. To avoid excessive migratory movement between fish from the up- and downstream sampling sites, the two sampling locations were chosen approximately 10 kilometers apart.

Six replicate water samples of 1 L were collected in sterile glass bottles, transported to the lab at 4°C, and vacuum-filtered through a 0.22 µm polycarbonate filter membrane (Ø 47 mm, Sartorius, Göttingen, Germany) until clogging on the same day. The total volume of filtered water ranged from 200 to 300 mL. Additionally, six replicate sediment samples per sampling site (20-50 cm depth) were collected in sterile 50 mL Falcon tubes. The filter membranes from the water samples and the sediment samples were stored at −20°C for subsequent DNA extraction.

Six fish of each species per sampling site - Brown trout (*Salmo trutta*) and European bullhead (*Cottus gobio*) - were sampled by electrofishing upstream and downstream of the WWTP. Fishing and animal handling were carried out in accordance with federal legislation and ethics approval based on permits issued by the Saxon State Office for Environment, Agriculture, and Geology (AZ 76/1/9222.22-03/22). Ethical aspects of sampling were conducted following the requirements of Directive 2010/63/EU of the European Parliament and of the Council of 22 September 2010 on the protection of animals used for scientific purposes [30]. Skin and gill samples were taken immediately after the fish were removed from the water using sterile cotton swabs. The collected samples were transferred into PowerBead tubes and stored at −20°C. The fish were then transported to the lab and stored at −20°C prior to dissection. Feces samples were obtained by dissecting the fish’s abdomen and extracting the intestinal contents. The feces were then transferred into a sterile tube and stored at −20°C before DNA extraction. Each fish sample was analyzed individually as a biological replicate.

### DNA extraction

DNA was extracted using the DNeasy PowerSoil Pro Kit (Qiagen, Hilden, Germany) according to the manufacturer’s protocol with some modifications depending on the sample type. The filter membrane was aseptically shredded into small fragments, transferred into PowerBead tubes, and used for DNA extraction. The sediment samples were centrifuged to remove excess water, homogenized, and 250 mg were weighed in for DNA extraction. DNA extraction from mucosal skin and gill filament samples was performed from cotton swabs. The fish feces were manually homogenized with a spatula, and then 250 mg of feces were used for DNA extraction. For the samples with less than 250 mg of feces, the entire available sample was extracted. The concentration of the extracted DNA was measured using NanoDrop 2000 (Thermo Fischer, Waltham, MA, USA) and the DNA was stored at −20°C for downstream analysis.

### DNA Sequencing

To analyze the bacterial community composition of the extracted DNA, samples were submitted to the Institute of Clinical Molecular Biology (IKMB, Kiel, Germany) for 16S rRNA gene-based amplicon sequencing. Sequencing was performed using the Illumina NovaSeq Platform with PCR primers targeting the V3-V4 region (V3F: 5′-CCTACGGGAGGCAGCAG-3′ V4R: 5′- GGACTACHVGGGTWTCTAAT-3) of the bacterial 16S rRNA gene [31]. Sequencing data was processed using the Mothur software package v.1.47.0 [32] as per the following procedure: Raw reads were merged using the ‘make.contigs’ command. Low-quality sequences were removed using the ‘screen.seqs’ command with maxambig = 0 and minlength = 300. Sequences were aligned to the reference SILVA 132 reference database [33] using the ‘align.seqs’ command and taxonomically classified against the RDP database [34] using the ‘classify.seqs’ command with cutoff = 80. Eukaryotic, chloroplast, archaeal, and mitochondrial sequences were removed using the ‘remove.lineag’ command. The chimera.vsearch command was used to identify and remove chimeric sequences. The ‘dist.seqs’ command was used to calculate pairwise sequence distances and was followed by the ‘cluster’ command to group sequences into OTUs based on the 97% similarity cutoff. To ensure equal sequencing depth across all samples, subsampling was performed using the ‘sub.sampl’ command in Mothur, normalizing the dataset to 3,791 reads per sample. Raw sequencing data was submitted to the NCBI Sequence Read Archive (SRA) under accession number PRJNA1158582.

### High throughput qPCR

To determine the relative abundance of individual target genes in the samples, isolated DNA samples were submitted to Resistomap Oy (Helsinki, Finnland) for high throughput qPCR using a SmartChip Real-time PCR system (TaKaRa Bio, Japan). The target genes included 27 ARGs and the 16S rRNA gene (Table S1) [35]. All samples were run with three technical replicates. The protocol was as follows: PCR reaction mixture (100 nL) was prepared using SmartChip TB Green Gene Expression Master Mix (TaKaRa Bio, Japan), nuclease-free PCR-grade water, 300 nM of each primer, and 2 ng/μL DNA template. After initial denaturation at 95 °C for 10 min, PCR comprised 40 cycles of 95 °C for 30 s and 60 °C for 30 s, followed by melting curve analysis for each primer set. A cycle threshold (Ct) of 27 was selected as the detection limit [36,37]. The quantification limit was calculated as 25 gene copies per reaction accounting for 12.5 gene copies per ng of DNA template. Amplicons with non-specific melting curves or multiple peaks were excluded. The relative abundances of the detected gene normalized to the 16S rRNA gene were estimated using the ΔCT method based on mean CTs of three technical replicates [38].

### Data processing and visualization

All data were analyzed in R.Studio v.2024.4.0.735 [39] and visualized using the R package ggplot2 v.3.4.3 [40]. Non-metric Multidimensional Scaling (NMDS) analysis was performed using the function *metaMDS* from the Vegan R package [41]. The Euclidean distance was used to calculate the distance matrix for ARGs, while the OTUs distance matrix was computed based on Bray-Curtis dissimilarities. Analysis of similarities (ANOSIM) tests was conducted using the *anosim* function from the Vegan package [41]. To test for differences in the abundance of ARGs or bacterial phyla between sampling locations two-tailed t-tests with Bonferroni correction for multiple testing were performed. Spearman’s rank correlation was used to test if significant differences in ARG abundance between up- and downstream samples can be explained by the abundance of the ARG in the WWTP effluent. Throughout, statistical significance was assumed for *P* < 0.05.

## Results

### Variations in bacterial community composition of water and sediment up- and downstream of the WWTP

To assess the effect of the WWTP effluent on the river water microbiome, we first compared the bacterial composition of water collected at the up- and downstream sampling sites. Across all water samples, the composition of bacterial communities was generally dominated by the phyla Proteobacteria, Actinobacteria, Bacteroidetes, and Firmicutes (Fig. 1A). Downstream of the WWTP, the bacterial composition exhibited a slight but significant increase in the relative abundance of the phyla Proteobacteria (up = 40.2 ± 2.0%, down = 57.7 ± 6.3%, *P* = 0.0009, two-tailed t-test) and Firmicutes (up = 3.5 ± 1.6%, down = 6.1 ± 2.2%, *P* = 0.046), as well as more than a five-fold significant increase in Verrucomicrobia (up = 0.9 ± 0.5%, down = 5.8 ± 3.5%, *P* = 0.019). Conversely, the river samples downstream exhibited a more than two-fold significant decrease in Actinobacteria (up = 13.2 ± 1.3%, down = 5.9 ± 2.1%, *P* = 0.0001) and a slight, non-significant decrease in the relative abundance of Bacteroidetes (up = 18.3 ± 3%, down = 12.3 ± 6.1%, *P* = 0.06). At the OTU level, differences between bacterial communities resulted in significantly distinct clusters of samples taken up- and downstream based on Bray-Curtis dissimilarity (R = 0.77, *P* = 0.005, ANOSIM; Fig. 1B).

**Figure 1.**
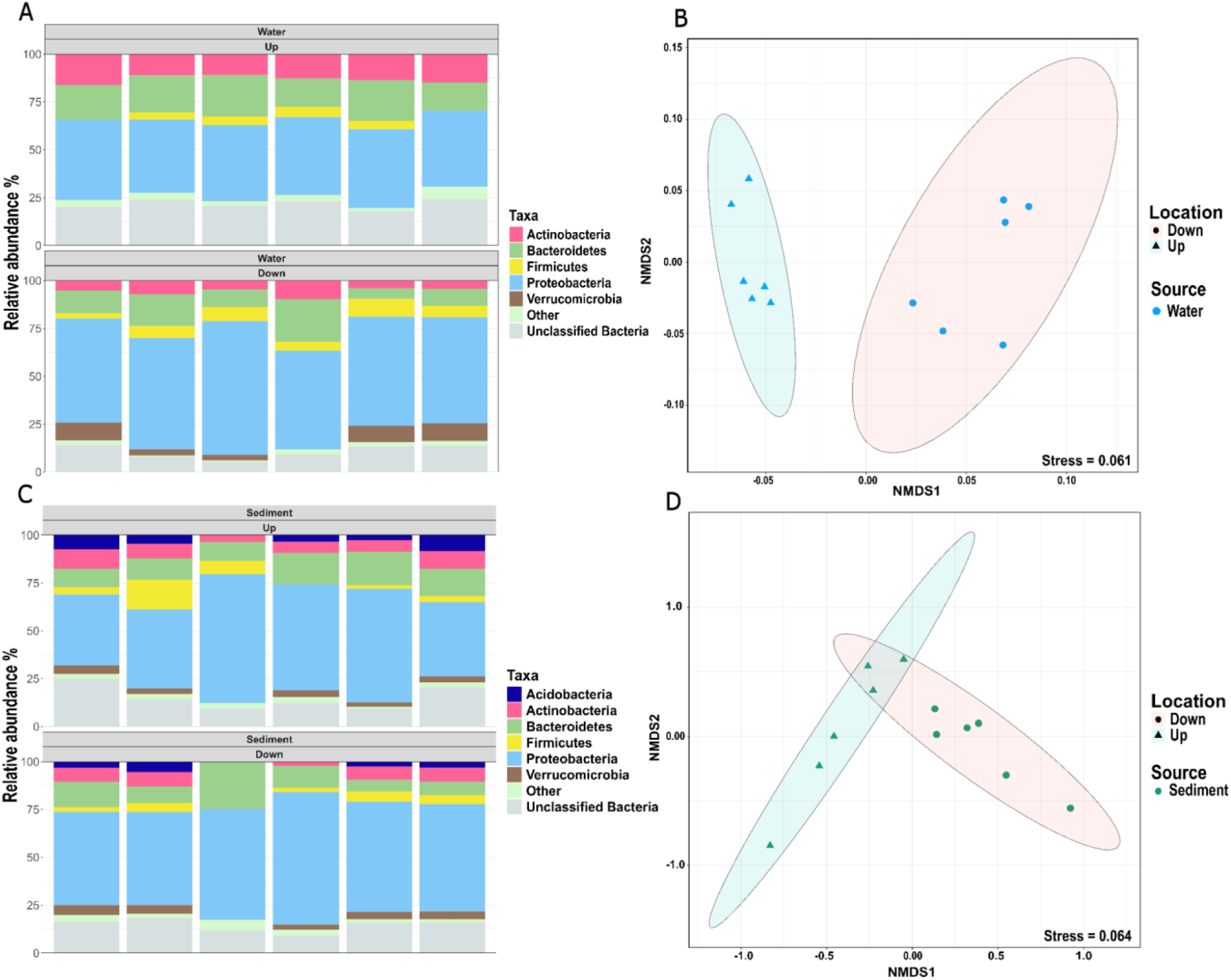
Bacterial community composition of river water and sediment samples up- and downstream of the WWTP. (A) Relative abundance of the dominant phyla present in river water samples. (B) Non-metric multidimensional scaling (NMDS) plot based on Bray-Curtis dissimilarities between water samples at the OTU level (97% sequence similarity). (C) Relative abundance of the dominant phyla present in river sediment samples. (D) Nonmetric multidimensional scaling (NMDS) plot based on Bray-Curtis dissimilarities between sediment samples at the OTU level (97% sequence similarity). Dominant phyla are defined as those with an average relative abundance of more than 2% across six replicate samples. The remaining phyla are grouped as “Other”. The ellipses indicate the 95% confidence intervals of grouping up- and downstream samples.

The river sediment bacterial communities were dominated by the same phyla as the river water. However, no significant differences between the sediment samples taken up- and downstream of the WWTP were observed for the dominant phyla: Proteobacteria (up = 49.9 ± 12.5%, down = 56.3 ± 7.5%, *P* = 0.3), Bacteroidetes (up = 13.2 ± 3.4%, down = 11.8 ± 6.8%, *P* = 0.66), Actinobacteria (up = 7 ± 2.5%, down = 5.5 ± 2.7%, *P* = 0.33), Acidobacteria (up = 4.5 ± 2.5%, down = 2.7 ± 1.5%, *P* = 0.2), Firmicutes (up = 5.5 ± 5.2%, down = 3.5 ± 1.8%, *P* = 0.4) and Verrucomicrobia (up = 2.7 ± 1.3%, down = 3.5 ± 1.5%, *P* = 0.2, Fig. 1C). Despite this, significant differences at the OTU level composition were still detected in the NMDS plot (Fig. 1D). However, the effect size of the ANOSIM test (*R* = 0.3713, *P* = 0.002) was far lower for sediment than for water (*R* = 0.77, *P* = 0.005, ANOSIM), indicating that WWTP effluent had a more pronounced effect on the aquatic rather than sediment fraction of the river. Still, both compartments of the studied river were clearly affected by WWTP effluents concerning microbial community composition at the OTU level.

### Microbiomes of trouts and bullheads vary between locations

To identify if the difference in the bacterial community composition between water and sediment samples was mirrored in alterations of the respective fish microbiomes, we investigated the bacterial composition of skin, gill, and fecal samples from trout and bullhead caught at the up- and downstream locations of the WWTP. Again, the phyla Proteobacteria, Actinobacteria, Firmicutes, and Bacteroidetes were predominant in all trout samples at both locations (Fig. 2A). Among them, the relative abundance of Verrucomicrobia was significantly higher downstream of the WWTP than upstream in all trout samples: in the skin (up = 0.26 ± 0.22%, down = 3.8 ± 2.3%, *P* = 0.01), gills (up = 0.72 ± 0.85%, down = 3 ± 1.4%, *P* = 0.009), and feces (up = 0.44 ± 0.34%, down = 1.6 ± 0.4%, *P* = 0.0006). This mirrored the effects observed for the river water where Verrucomicrobia displayed the highest increase by up to five-fold in downstream communities. The decrease in Bacteroidetes and Actinobacteria observed in the downstream river water samples was also observed in trout skin with a two-fold significant decrease in Bacteroidetes (up = 13 ± 4.1%, down = 6.6 ± 2.4%, *P* = 0.01) and Actinobacteria (up = 10.7 ± 3.8%, down = 4.5 ± 1.3%, *P* = 0.008). The relative abundance of other dominant phyla did not exhibit statistically significant differences between both locations.

**Figure 2.**
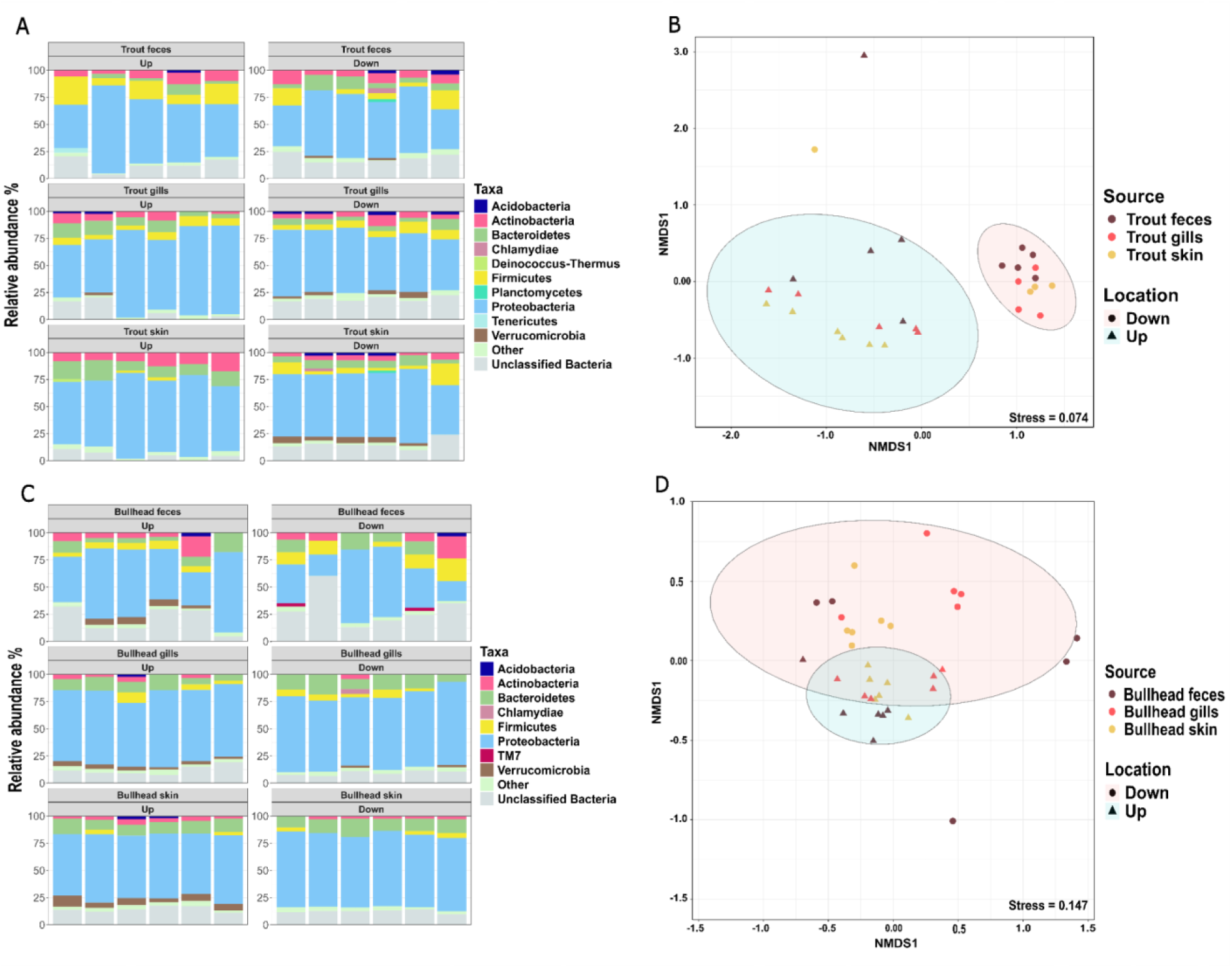
Bacterial community composition of the trout and bullhead samples up- and downstream of the WWTP. (A) Relative abundance of the dominant phyla present in trout samples. (B) Non-metric multidimensional scaling (NMDS) plot based on Bray-Curtis dissimilarities between trout samples at the OTU level (97% sequence similarity). No significant grouping based on the three different trout sample types was observed. (C) Relative abundance of the dominant phyla present in bullhead samples. (D) Nonmetric multidimensional scaling (NMDS) plot based on Bray-Curtis dissimilarities between bullhead samples at the OTU level (97% sequence similarity). No significant grouping based on the three different bullhead sample types was observed. Dominant phyla are defined as those with an average relative abundance of more than 2% across six replicate samples. The remaining phyla are grouped as “Other”. The ellipses indicate the 95% confidence intervals of grouping up- and downstream samples.

Comparison of the bacterial communities between the trout samples up- and downstream of the WWTP at the OTU level resulted in two distinct clusters, confirming the difference in microbial community composition of the trout samples (*R* = 0.7733, *P* = 0.001, ANOSIM). Interestingly, no significant grouping based on the three different sample types was observed within the clusters, indicating the sample types’ homogeneity (Fig. 2B).

In the case of bullhead, the phyla Proteobacteria, Bacteroidetes, Firmicutes, Verrucomicrobia, and Actinobacteria were also most prevalent in the microbial communities at both locations. However, the difference between the locations was less pronounced. This aligns with a higher similarity of dominant phyla observed in the sediment samples, as bullheads are rather sediment-associated. A notable difference between the locations was the significant increase in Proteobacteria levels in the skin samples collected downstream of the WWTP, where they reached 67.6 ± 1.7% compared to 59.3 ± 3.2% upstream (*P* = 0.0006, Fig. 2C).

The low degree of dissimilarity of the bullhead microbiomes up- and downstream of the WWTP was supported by the visual inspection of the NMDS plot with the ellipses largely overlapping (Fig. 2D). At the same time, the conducted ANOSIM still revealed a significant dissimilarity between the locations, but the effect size was only moderate compared to trout or water samples (*R*=0.23, *P* = 0.001).

An observable difference between the microbiomes of the two different fish was that the trout microbiomes consolidated from a more diverse, variable microbiome upstream towards a more similar and consistent microbiome downstream of the WWTP (Fig. 2B). Contrary, the bullhead microbiomes displayed the opposite trend (Fig. 2D). This could, in theory, be connected to their difference in lifestyle, e.g., being mainly exposed to the water (trout) or sediment (bullhead) microbiome. If this is the case, the diversity of the pool of microbes available for the fish microbiome to recruit from might be affected downstream of the WWTP in these different environmental compartments. However, no significant differences in Shannon diversity of either the water (up = 5.08 ± 0.29, down = 4.59 ± 0.29, *P* = 0.4517, two-tailed t-test) or the sediment microbiome (up = 5.77 ± 0.72, down = 5.48 ± 0.56, *P* = 0.4523) was detected (Fig. S2).

### Water and sediment exhibit low variation in ARG profiles between locations

To investigate the potential effect of WWTP effluent on the resistomes of river water and sediment, we next compared the resistomes of the environmental samples collected upstream and downstream of the WWTP. The difference in ARG composition in water samples between the two locations resulted in two distinct clusters on the NMDS plot (Fig. 3A). The ANOSIM statistical test indicates moderate dissimilarity between these two groups (*R* = 0.2981, *P* = 0.018), which partially reflects the differences in the water microbiomes. In contrast, the comparison of sediment samples taken up- and downstream of the WWTP revealed strong similarities in sediment ARG profiles across locations, as confirmed by the ANOSIM test (*R* = −0.0277, *P* = 0.58) (Fig. 3B). After we observed overlapping yet distinct clusters of the water resistomes up- and downstream of the WWTP on the NMDS plot, indicating no general response on all ARGs, we next focused on identifying those ARGs for which the relative abundance downstream was increased by WWTP effluent.

**Figure 3.**
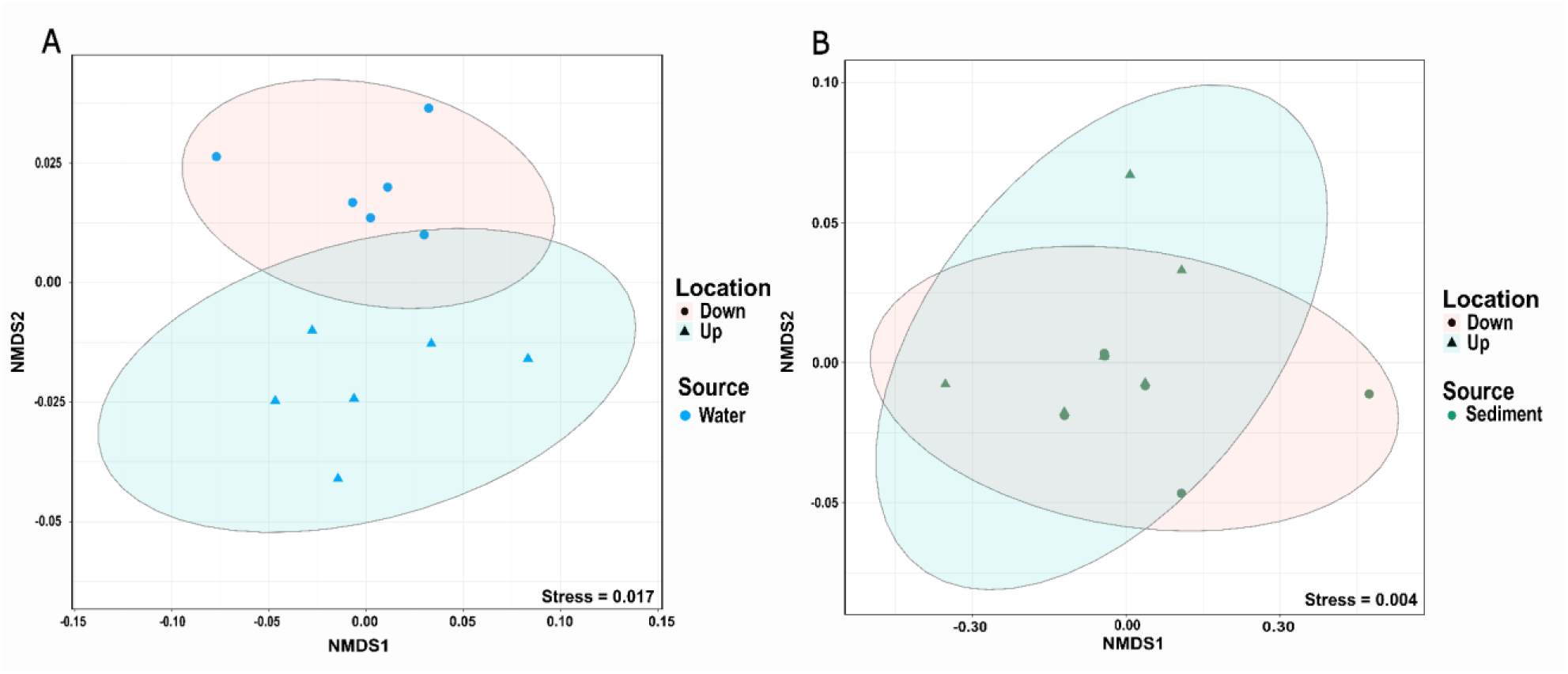
NMDS plot of ARGs composition of (A) water and (B) sediment samples up- and downstream the WWTP. The distance matrix was computed based on the Euclidean distance. The radius of the ellipses represents the 95% confidence intervals of grouping up- and downstream samples.

### WWTP effluent increases the level of specific individual ARGs downstream of the WWTP

To identify ARGs whose relative abundance was impacted by the WWTP effluent, we first compared the relative abundances of the tested ARGs up- and downstream of the WWTP. We aimed for the ARGs whose relative abundance was higher downstream than upstream of the WWTP and for which the difference in the relative abundance was statistically significant. Furthermore, to pinpoint that such increases could indeed stem from WWTP effluents we analyzed if these ARGs were significantly more prevalent in WWTP effluent by calculating the ratio of the relative abundances of the ARGs detected in the WWTP effluent compared to the relative abundance observed upstream. In total, we identified eight ARGs that were significantly increased in relative abundance downstream of the WWTP (*P* < 0.05, two-tailed t-test with Bonferroni correction for multiple testing). Out of these, six were at least two-fold higher in relative abundance in the effluent compared to the upstream samples: *bla*_OXA-58_*, ermB*, *ermF, tetW, vanA,* and *sul1* (Fig. 4). Moreover, all four ARGs with highly increased abundance by more than 10-fold in wastewater effluent compared to upstream samples displayed a significantly higher abundance in downstream samples further indicating a clear effect of WWTP effluent on the water resistomes at the individual ARG level. Statistically, the likelihood of an ARG to be significantly more abundant downstream positively correlated with the fold-change in abundance between upstream and WWTP effluent samples (*P* = 0.00172, *r_S_* = 0.58434, Spearman’s rank correlation, Fig. 4).

**Figure 4.**
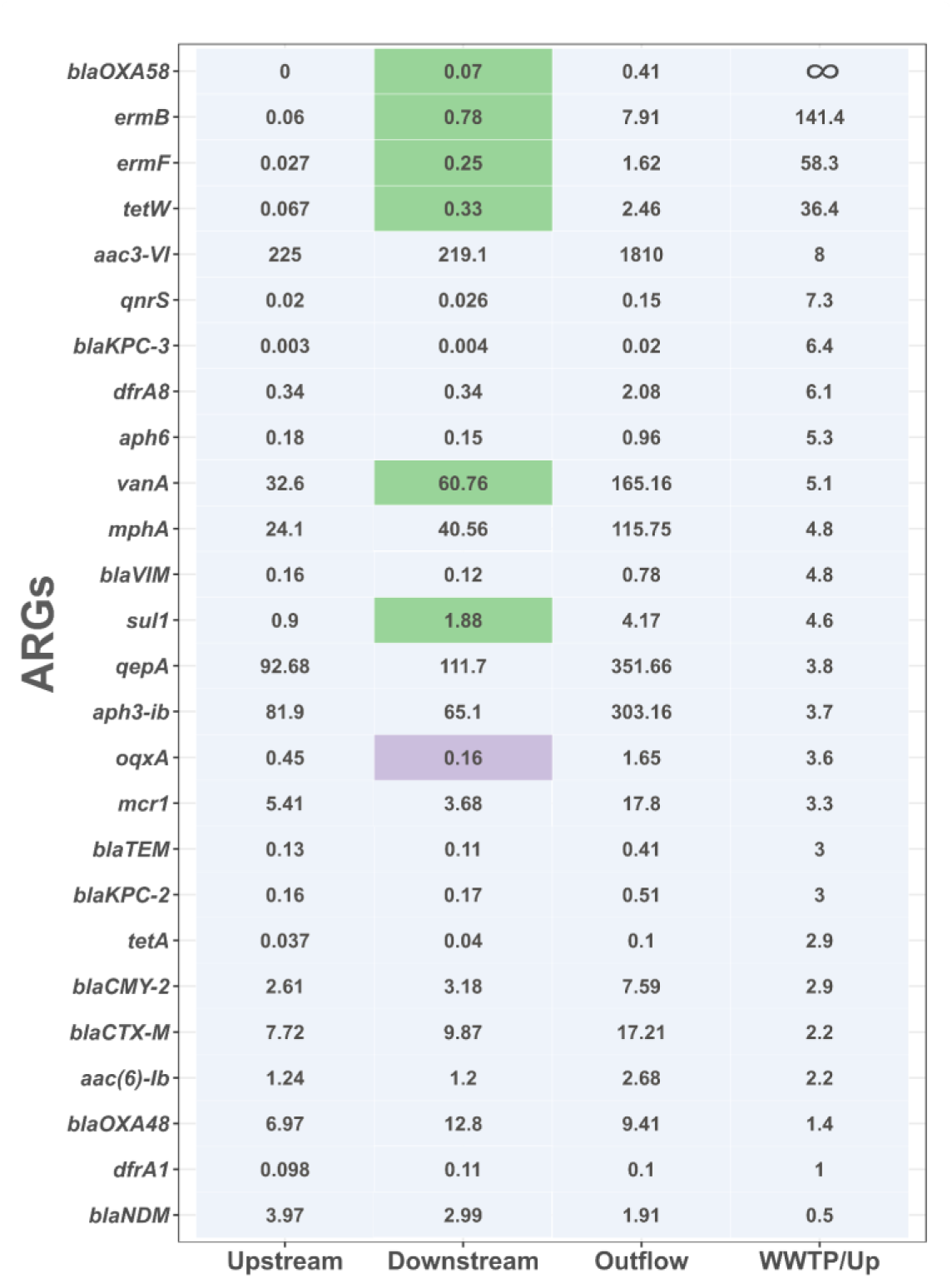
Normalized relative abundance of the ARGs per 1000 copies of the bacterial 16S rRNA gene in water samples. The numbers represent the average relative abundance of six biological replicates. The WWTP/Upstream ratio was calculated by dividing the average relative abundance of each gene upstream of the WWTP by the relative abundance of the WWTP effluent. ARGs are presented in descending order of their P >WWTP/Upstream ratio. Ratios, where the ARG was not detected upstream but was detected in the WWTP effluent, are given as ∞. A two-tailed t-test with Bonferroni correction for multiple testing was performed to test the statistical significance of the difference in relative abundance of ARGs up- and downstream of the WWTP. Relative abundance highlighted in green indicates those ARGs with a statistically significant increase of the relative abundance downstream of the WWTP (*P*<0.05). Relative abundance highlighted in purple shows the ARGs with a statistically significant decrease in the relative abundance of the ARGs (P<0.05). More detailed values, including standard deviations, can be found in Figure S3.

### ARG composition in fish samples remains consistent across locations

To investigate whether these differences observed in water ARG profiles up- and downstream of the WWTP could also be seen in fish, we compared the resistomes of the trout and bullhead samples up- and downstream of the WWTP. Although the dissimilarity of the water microbiomes was mirrored in trout microbiomes, the resistomes of trout were indistinguishable at both sites (*R* = 0.1468, *P* = 0.21, ANOSIM). Dissimilarities of bullhead samples between the two locations were while appearing significant only having a very low effect size (*R* = 0.08, *P* = 0.034, ANOSIM), which is consistent with the low degree of difference observed between bullhead microbiomes obtained up- and downstream (Fig. 5). When compared individually, no significant differences were found between the gills, skin, or feces of both fish species up- and downstream of the WWTP (all *P* > 0.05, ANOSIM). Since we did not observe a general impact of the wastewater effluent on the fish resistomes, we further investigated if the wastewater effluent affects the level of individual ARGs in fish samples similar to what was observed for water samples.

**Figure 5.**
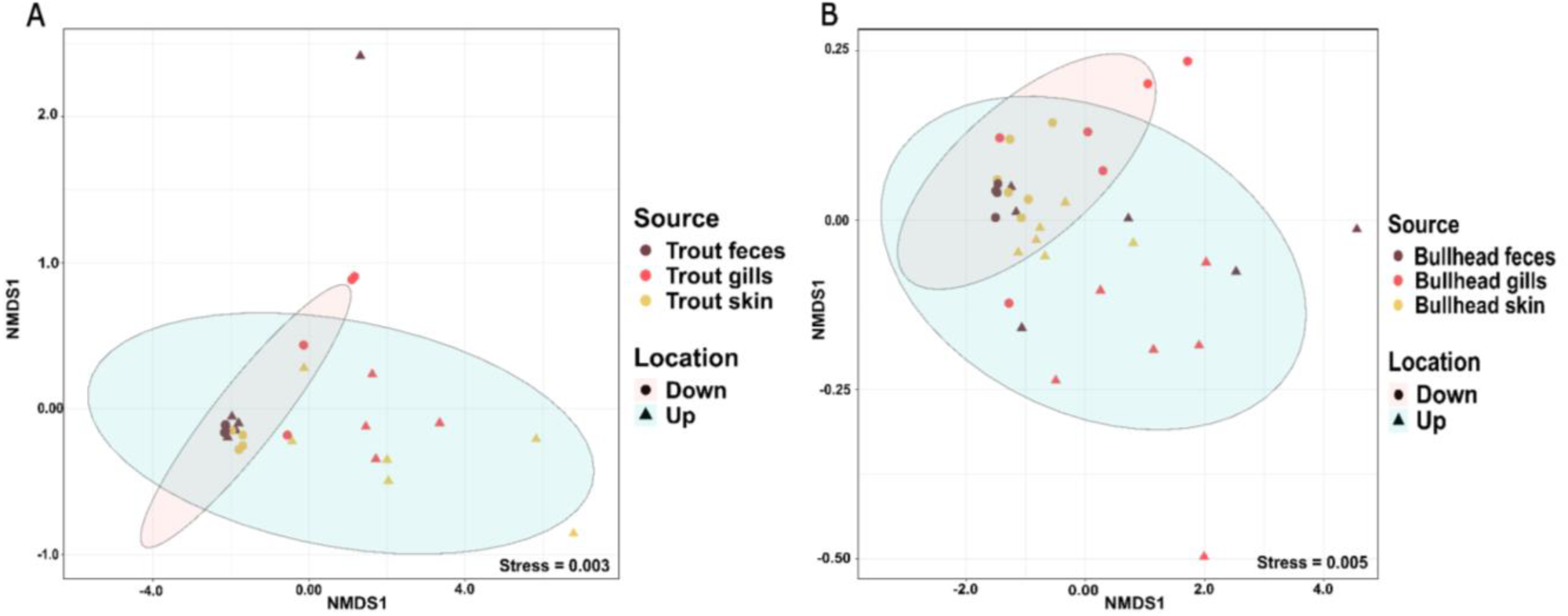
NMDS plot of ARG composition of (A) trout and (B) bullhead samples up- and downstream the WWTP. The distance matrix was computed based on the Euclidean distance. The radius of the ellipses represents a 95% confidence interval of grouping up- and downstream trout and bullhead samples.

### Wastewater effluent effects on ARGs in water samples are not mirrored in fish

After we observed that WWTP effluent increased the abundance levels of at least seven tested ARGs, we considered these genes as potential indicators for the evaluation of trout and bullhead as effective sentinels for AMR pollution: we tested if the difference in the relative abundance of these ARGs in water corresponded to those in trout and bullhead.

For that, we first compared the relative abundance of the ARGs in the trout samples up- and downstream of the WWTP. None of the tested ARGs exhibited significantly higher abundance downstream of the WWTP (Fig. 6). We observed similar results when comparing ARG levels in bullhead samples from upstream and downstream locations (Fig. S5). Hence, the resistomes of both trout and bullhead did not reflect the changes in the water resistome caused by wastewater effluent. This was observed both in terms of overall resistome effects and at the level of individual ARGs.

**Figure 6.**
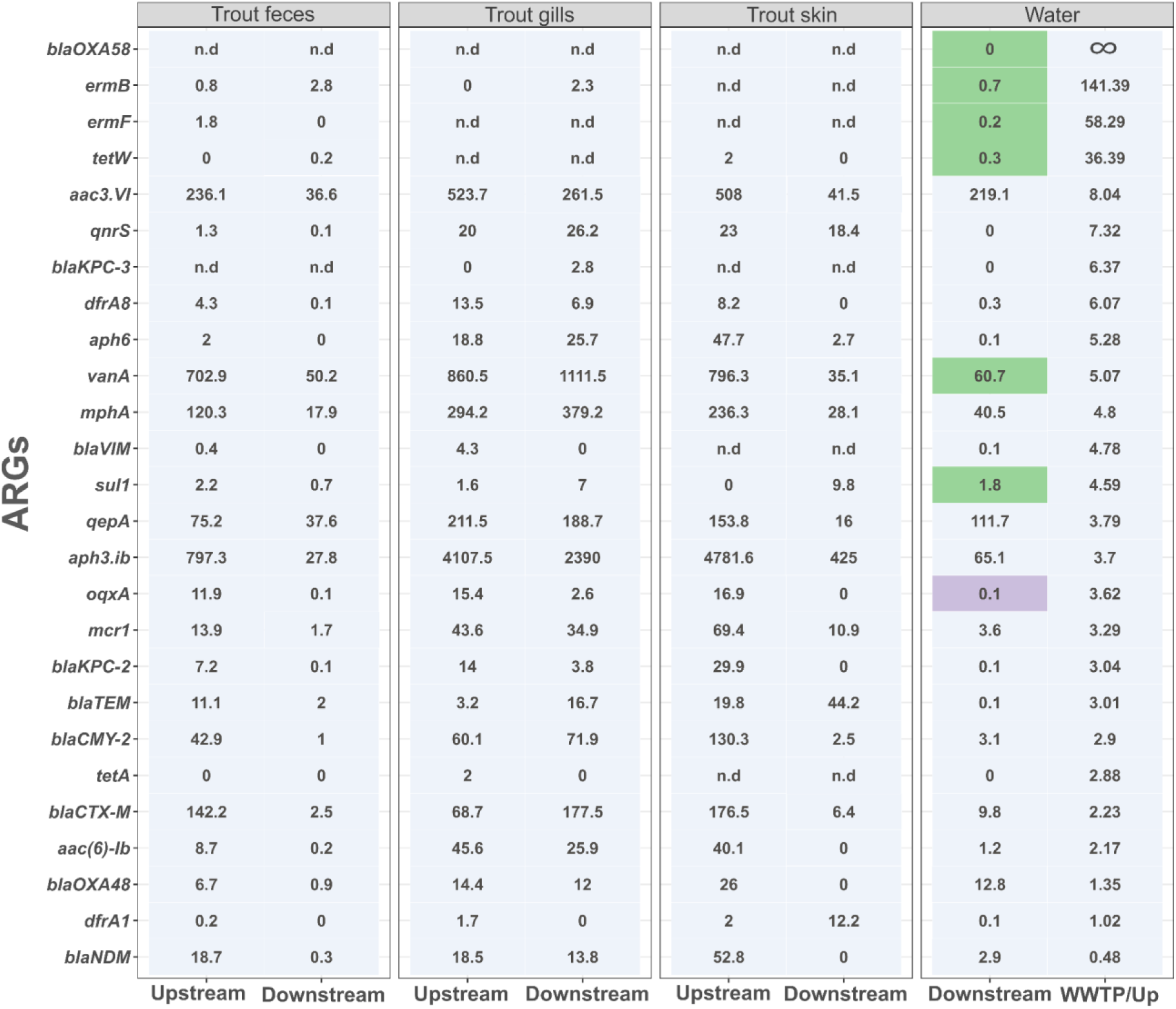
Normalized relative abundance of the ARGs per 1000 copies of the bacterial 16S gene in trout samples. The numbers represent the average relative abundance of the biological replicates. A two-tailed t-test with Bonferroni correction for multiple testing was performed to test the statistical significance of the difference in relative abundance of ARGs up- and downstream of the WWTP. Relative abundance highlighted in green indicates the ARGs with a statistically significant increase of the relative abundance downstream of the WWTP (P<0.05). Relative abundance highlighted in purple is the ARGs with a statistically significant decrease in the relative abundance of the ARGs (P<0.05). The column with the relative abundance of the ARGs observed in water downstream of the WWTP indicates the ARGs, whose increase in relative abundance was impacted by the effluent. Ratios, where the ARG was not detected in water samples upstream of the WWTP but was detected in the WWTP effluent, are given as ∞. More detailed values, including standard deviations, can be found in Figure S4.

## Discussion

In this study, we investigated whether two freshwater fish species - Brown trout and European bullhead - are potential sentinels for AMR pollution in rivers receiving effluents from municipal WWTPs. Wastewater effluent had a pronounced effect on the microbiome and resistomes of the riverine water and to a lesser degree on the riverine sediment downstream of the WWTP. The microbiomes of both fish species reflected the differences in bacterial composition observed in water. However, the resistome compositions of the trout and bullhead samples displayed barely any dissimilarity between locations, despite the effects of WWTP effluent observed for the resistome in the water samples.

The observed effects of WWTP effluent leading to altered river microbiomes with higher abundances of ARGs downstream of the WWTP are in line with numerous previous reports [18–21,42]. In this study, the distance to the WWTP (3.6 km downstream) was higher than in most comparable studies. Still, clear effects on the microbiome and the resistome of the river water were observed. We further provide strong support that these differences between up- and downstream microbiomes and resistomes in the water are indeed caused by the WWTP effluent despite the distance and not only local variation. First, the likelihood of an ARG to be significantly more abundant downstream positively correlated with the fold-change in abundance between upstream and WWTP effluent samples. In addition, the river samples collected downstream of the WWTP displayed increased levels of bacteria belonging to the phyla Proteobacteria and Firmicutes, both of which were most abundant in wastewater effluent samples and are typical for general wastewater bacterial communities [43]. Although several studies have reported a strong long-term impact of WWTP effluent on sediment bacterial communities and resistomes [44–47], mainly because both abiotic pollutants as well as ARB can settle and accumulate at the river ground. However, we observed a weaker effect on the sediment in this study. This can be explained by sampling sites in previous studies being regularly located close to the WWTP (0.05 to 1.5 kilometers) where the majority of settling of particles takes place [45–47]. The greater distance to the WWTP discharge point in our study (>3 km) where most particles already have settled and accumulated in the sediment before the sampling point may explain why the effect of the effluent was here more pronounced in the water column than the sediment.

After establishing that WWTP effluent affected the downstream water microbiome, we here report that the effects are largely mirrored in the fish microbiomes. This was particularly evident for the bacterial composition of fish gill and skin microbiomes, which are susceptible to environmental impact and often depend on the water microbiome due to their direct and constant exposure [48,49]. Similarly, fish feces microbiomes cluster distinctly up- and downstream of the WWTP but together with the respective gill and skin microbiomes. This can be attributed to the significant influence of the constantly ingested water on the gut microbiome [49,50]. Throughout, the same dominant phyla Proteobacteria, Bacteroidetes, Actinobacteria, and Firmicutes were observed in all fish samples, with these phyla being common members of freshwater fish microbiomes [48,51,52]. The homogeneity in bacterial composition and consistency of effects of the WWTP effluent across all fish sample types and for either fish species clearly displays the significant influence of the surrounding river environment on the fish microbiome.

Still, clear effects of the lifestyle and behavior of the two different fish species became apparent when analyzing the strength of the WWTP effluent effects on the microbiomes. The far higher degree of dissimilarity between up- and downstream locations observed for the trout samples aligns with the more pronounced effects the WWTP effluent had on the water compared to the sediment samples. Trout typically actively live in the water column [25,26], meanwhile, bullheads are benthic, sedentary, sediment-associated fish [27,28]. Hence, the moderate dissimilarities in the bullhead microbiome are reflective of the moderate but significant differences in the sediment up- and downstream.

Contrarily, the effects of WWTP effluent on the water and sediment overall resistome composition and individual ARG abundances were not reflected in the fish resistomes downstream of the WWTP. No effects being observed for the benthic bullhead resistome can likely be explained by its sediment-associated lifestyle [27,28], as the effects of WWTP effluent on the sediment resistome were, while significant, rather small. A larger effect would have been expected for the free-water associated trout [25,26] as far more significant effects were observed for the water resistome. However, similar to the bullhead, high similarity of the up- and downstream resistome was observed with no clear effects on overall resistome composition or individual ARG abundances. This high resistome homogeneity suggests that the ARGs detected in the fish samples are rather associated with the natural fish commensal microbiota and not introduced from the external environment. Previous reports confirm that the natural fish microbiome can indeed host a plethora of diverse ARGs [53,54]. When analyzing the resistome in isolation, a second plausible explanation is that fish are dynamic living organisms that can travel from upstream to downstream of the WWTP in the river. Hence the similarities in the resistome could originate from being exposed at both locations. However, this should also be reflected in the fish microbiome exhibiting limited local variation. Contrary to this, we observed significant differences in the microbiomes up- and downstream for either of the two tested fish species, with higher microbiome dissimilarities in the even more active swimmer trout [25,26] being particularly remarkable. Consequently, fish movement between sampling locations can likely be excluded as a main contributing factor.

In the frameworks of environmental surveillance [4,7,55,56] and subsequent human health-based risk assessment of environmental AMR [27,57–59] crucial factors include ARG hosts, mobility, likelihood of human exposure, and the ability to invade and colonize the human microbiome. The colonization potential of humans with ARB through environmental exposure was already highlighted by a previous study documenting that surfers who regularly swallow increased amounts of seawater (often affected by WWTP effluents) are indeed more likely colonized by AMR *E. coli* compared to a non-surfing control group [60]. To include such colonization risks in the environmental surveillance of AMR, several studies have explored the potential of wildlife species to serve as sentinels for the colonization potential connected to environmental AMR pollution. For example, wild, small mammals living near water bodies receiving effluent from WWTPs are more likely to carry antibiotic-resistant *E. coli* than those residing farther from polluted rivers [61,62]. Similarly, seagulls frequenting WWTPs and landfills were increasingly colonized by pathogenic ARBs [63]. However, we here demonstrate that using fish as sentinels is less promising, likely as the fish gut, skin and gill environment is rather selective for bacterial species that are less receptive to host WWTP borne ARGs. We established that the fish microbiome is not resilient to the changes in the river microbiome through WWTP effluent in general, for example, the shift in relative abundance of the phylum Verrucomicrobia in river water was reflected in trout. However, the observed microbiome shifts did not significantly affect the ARG composition in either trouts or bullheads downstream of the WWTP. This is likely due to a strong association of the tested ARGs with specific host ARB. Indeed previous reports indicate that the resistome of WWTP effluent strongly and significantly correlates with its microbiome composition [64,65], highlighting the importance of specific ARG-host associations. Hence, it is not the overall colonization potential of the environmental microbiomes that is relevant here, but rather the colonization potential of specific ARB carrying the clinically relevant ARGs being tested. Within this context, it needs to be considered that a majority of bacteria in municipal wastewater effluents, and particularly the proportion carrying the clinically relevant ARGs, typically consist of bacterial species that originated from human or other warm-blooded animal gut microbiomes [11,17]. The resilience of the fish resistome towards colonization by these effluent-associated ARBs could hence be linked to these ARBs being maladapted to colonization under conditions different than the digestive tract of warm-blooded animals. Therefore, the emergence of ARB and ARG detected in the wastewater effluent in poikilothermic organisms, such as fish is less probable. This notion is further supported as effects of WWTP effluent in Swiss rivers were evident in the water resistome, while no significant shift in the resistome composition of poikilothermic river amphipods was detected [19]. Meanwhile, animals with physiological characteristics similar to humans, such as small mammals [61,62] or birds [63], and with high proportions of Proteobacteria, as particularly prominent ARG hosts and recipients, in their gut microbiome [66] are more susceptible to colonization by ARB and ARG originating from wastewater effluent. This demonstrates that using the colonization potential of wildlife as sentinels can become an effective mechanism for evaluating AMR-associated risks as part of the surveillance of key environmental AMR hotspots, such as WWTPs and their surroundings [61,62]. However, our results indicate that not all wildlife species are equally suitable for such AMR surveillance efforts and that fish, despite having a high exposure in WWTP effluent-affected waters, are not well suited as sentinels for assessing AMR colonization risks.

## Acknowledgments

The authors thank Felix Grunicke and Luise Richter for support with sampling and fish handling and Steffen Kunze, Christiane Zschornack, and Juliane Isler for technical support in the laboratory. The authors further thank the EMBARK and SEARCHER consortia for discussion and improving the interpretations presented in this manuscript.

## Data Availability

The datasets supporting the conclusions of this article are included within the article and its additional files or available through the corresponding author upon reasonable request. Raw data regarding the high-throughput qPCR assays is available in Supplementary Table 2. Original sequencing data is available in the NCBI sequencing read archive under project accession number PRJNA1158582 with individual sample accession codes given in Supplementary Table 3.

## Author Contributions - CRediT

**Faina Tskhay:** Conceptualization; Methodology; Validation; Formal analysis; Investigation; Data Curation; Writing - Original Draft; Writing - Review & Editing; Visualization

**Christoph Köbsch:** Conceptualization; Methodology; Writing - Review & Editing; Project administration

**Alan X. Elena:** Methodology; Formal analysis; Data Curation; Writing - Review & Editing

**Johan Bengtsson-Palme:** Conceptualization; Writing - Review & Editing; Project administration

**Thomas U. Berendonk:** Conceptualization; Resources; Writing - Review & Editing; Visualization; Supervision; Project administration; Funding acquisition

**Uli Klümper:** Conceptualization; Methodology; Resources; Data Curation; Writing - Original Draft; Writing - Review & Editing; Visualization; Supervision; Project administration; Funding acquisition

## References

[1] World Health Organization, A One Health Priority Research Agenda for Antimicrobial Resistance, Food & Agriculture Org., 2023.

[2] M.E. Velazquez-Meza, M. Galarde-López, B. Carrillo-Quiróz, C.M. Alpuche-Aranda, Antimicrobial resistance: One Health approach, Vet. World 15 (2022) 743–749. 10.14202/vetworld.2022.743-749.

[3] J. Bengtsson-Palme, E. Kristiansson, D.G.J. Larsson, Environmental factors influencing the development and spread of antibiotic resistance, FEMS Microbiol. Rev. 42 (2018). 10.1093/femsre/fux053.

[4] T.U. Berendonk, C.M. Manaia, C. Merlin, D. Fatta-Kassinos, E. Cytryn, F. Walsh, H. Bürgmann, H. Sørum, M. Norström, M.-N. Pons, N. Kreuzinger, P. Huovinen, S. Stefani, T. Schwartz, V. Kisand, F. Baquero, J.L. Martinez, Tackling antibiotic resistance: the environmental framework, Nat. Rev. Microbiol. 13 (2015) 310–317. 10.1038/nrmicro3439.

[5] D.G.J. Larsson, W.H. Gaze, R. Laxminarayan, E. Topp, AMR, One Health and the environment, Nat. Microbiol. 8 (2023) 754–755. 10.1038/s41564-023-01351-9.

[6] D.G.J. Larsson, C.-F. Flach, Antibiotic resistance in the environment, Nat. Rev. Microbiol. 20 (2022) 257–269. 10.1038/s41579-021-00649-x.

[7] J. Bengtsson-Palme, A. Abramova, T.U. Berendonk, L.P. Coelho, S.K. Forslund, R. Gschwind, A. Heikinheimo, V.H. Jarquín-Díaz, A.A. Khan, U. Klümper, U. Löber, M. Nekoro, A.D. Osińska, S. Ugarcina Perovic, T. Pitkänen, E.K. Rødland, E. Ruppé, Y. Wasteson, A.L. Wester, R. Zahra, Towards monitoring of antimicrobial resistance in the environment: For what reasons, how to implement it, and what are the data needs?, Environ. Int. 178 (2023) 108089. 10.1016/j.envint.2023.108089.

[8] A. Pruden, P.J. Vikesland, B.C. Davis, A.M. de Roda Husman, Seizing the moment: now is the time for integrated global surveillance of antimicrobial resistance in wastewater environments, Curr. Opin. Microbiol. 64 (2021) 91–99. 10.1016/j.mib.2021.09.013.

[9] N. Czekalski, E. Gascón Díez, H. Bürgmann, Wastewater as a point source of antibiotic-resistance genes in the sediment of a freshwater lake, ISME J. 8 (2014) 1381–1390. 10.1038/ismej.2014.8.

[10] I. Michael, L. Rizzo, C.S. McArdell, C.M. Manaia, C. Merlin, T. Schwartz, C. Dagot, D. Fatta-Kassinos, Urban wastewater treatment plants as hotspots for the release of antibiotics in the environment: A review, Water Res. 47 (2013) 957–995. 10.1016/j.watres.2012.11.027.

[11] L. Rizzo, C. Manaia, C. Merlin, T. Schwartz, C. Dagot, M.C. Ploy, I. Michael, D. Fatta-Kassinos, Urban wastewater treatment plants as hotspots for antibiotic resistant bacteria and genes spread into the environment: A review, Sci. Total Environ. 447 (2013) 345–360. 10.1016/j.scitotenv.2013.01.032.

[12] D. Cacace, D. Fatta-Kassinos, C.M. Manaia, E. Cytryn, N. Kreuzinger, L. Rizzo, P. Karaolia, T. Schwartz, J. Alexander, C. Merlin, H. Garelick, H. Schmitt, D. de Vries, C.U. Schwermer, S. Meric, C.B. Ozkal, M.-N. Pons, D. Kneis, T.U. Berendonk, Antibiotic resistance genes in treated wastewater and in the receiving water bodies: A pan-European survey of urban settings, Water Res. 162 (2019) 320–330. 10.1016/j.watres.2019.06.039.

[13] A. Karkman, T.T. Do, F. Walsh, M.P.J. Virta, Antibiotic-Resistance Genes in Waste Water, Trends Microbiol. 26 (2018) 220–228. 10.1016/j.tim.2017.09.005.

[14] M. Samer, Wastewater Treatment Engineering, BoD – Books on Demand, 2015.

[15] H. Bürgmann, D. Frigon, W. H Gaze, C. M Manaia, A. Pruden, A.C. Singer, B. F Smets, T. Zhang, Water and sanitation: an essential battlefront in the war on antimicrobial resistance, FEMS Microbiol. Ecol. 94 (2018) fiy101. 10.1093/femsec/fiy101.

[16] W. Zieliński, E. Korzeniewska, M. Harnisz, J. Drzymała, E. Felis, S. Bajkacz, Wastewater treatment plants as a reservoir of integrase and antibiotic resistance genes – An epidemiological threat to workers and environment, Environ. Int. 156 (2021) 106641. 10.1016/j.envint.2021.106641.

[17] A. Novo, C.M. Manaia, Factors influencing antibiotic resistance burden in municipal wastewater treatment plants, Appl. Microbiol. Biotechnol. 87 (2010) 1157–1166. 10.1007/s00253-010-2583-6.

[18] J. Alexander, N. Hembach, T. Schwartz, Evaluation of antibiotic resistance dissemination by wastewater treatment plant effluents with different catchment areas in Germany, Sci. Rep. 10 (2020) 8952. 10.1038/s41598-020-65635-4.

[19] J. Lee, F. Ju, K. Beck, H. Bürgmann, Differential effects of wastewater treatment plant effluents on the antibiotic resistomes of diverse river habitats, ISME J. 17 (2023) 1993–2002. 10.1038/s41396-023-01506-w.

[20] A. Osińska, E. Korzeniewska, M. Harnisz, E. Felis, S. Bajkacz, P. Jachimowicz, S. Niestępski, I. Konopka, Small-scale wastewater treatment plants as a source of the dissemination of antibiotic resistance genes in the aquatic environment, J. Hazard. Mater. 381 (2020) 121221. 10.1016/j.jhazmat.2019.121221.

[21] S. Rodriguez-Mozaz, S. Chamorro, E. Marti, B. Huerta, M. Gros, A. Sànchez-Melsió, C.M. Borrego, D. Barceló, J.L. Balcázar, Occurrence of antibiotics and antibiotic resistance genes in hospital and urban wastewaters and their impact on the receiving river, Water Res. 69 (2015) 234–242. 10.1016/j.watres.2014.11.021.

[22] E. Kristiansson, J. Fick, A. Janzon, R. Grabic, C. Rutgersson, B. Weijdegård, H. Söderström, D.G.J. Larsson, Pyrosequencing of Antibiotic-Contaminated River Sediments Reveals High Levels of Resistance and Gene Transfer Elements, PLOS ONE 6 (2011) e17038. 10.1371/journal.pone.0017038.

[23] O. Barraud, L. Laval, L. Le Devendec, E. Larvor, C. Chauvin, E. Jouy, S. Le Bouquin, Y. Vanrobaeys, B. Thuillier, B. Lamy, S. Baron, Integrons from *Aeromonas* isolates collected from fish: A global indicator of antimicrobial resistance and anthropic pollution, Aquaculture 576 (2023) 739768. 10.1016/j.aquaculture.2023.739768.

[24] H. Abgottspon, M.T. Nüesch-Inderbinen, K. Zurfluh, D. Althaus, H. Hächler, R. Stephan, Enterobacteriaceae with Extended-Spectrum-and pAmpC-Type β-Lactamase-Encoding Genes Isolated from Freshwater Fish from Two Lakes in Switzerland, Antimicrob. Agents Chemother. 58 (2014) 2482–2484. 10.1128/aac.02689-13.

[25] E. Aparicio, R. Rocaspana, A. de Sostoa, A. Palau-Ibars, C. Alcaraz, Movements and dispersal of brown trout (Salmo trutta Linnaeus, 1758) in Mediterranean streams: influence of habitat and biotic factors, PeerJ 6 (2018) e5730. 10.7717/peerj.5730.

[26] D.F. Clapp, R.D. Clark Jr., J.S. Diana, Range, Activity, and Habitat of Large, Free-Ranging Brown Trout in a Michigan Stream, Trans. Am. Fish. Soc. 119 (1990) 1022–1034. 10.1577/1548-8659(1990)119<1022:RAAHOL>2.3.CO;2.

[27] G. Knaepkens, L. Bruyndoncx, M. Eens, Assessment of residency and movement of the endangered bullhead (Cottus gobio) in two Flemish rivers, Ecol. Freshw. Fish 13 (2004) 317–322. 10.1111/j.1600-0633.2004.00065.x.

[28] S. Roje, B. Drozd, L. Richter, J. Kubec, Z. Polívka, S. Worischka, M. Buřič, Comparison of Behavior and Space Use of the European Bullhead Cottus gobio and the Round Goby Neogobius melanostomus in a Simulated Natural Habitat, Biology 10 (2021) 821. 10.3390/biology10090821.

[29] EUR-Lex - 31992L0043 - EN, Off. J. 206 22071992 P 0007 - 0050 Finn. Spec. Ed. Chapter 15 Vol. 11 P 0114 Swed. Spec. Ed. Chapter 15 Vol. 11 P 0114 (n.d.). https://eur-lex.europa.eu/legal-content/EN/TXT/HTML/?uri=CELEX%3A31992L0043 (accessed August 27, 2024).

[30] Directive - 2010/63 - EN - EUR-Lex, (n.d.). https://eur-lex.europa.eu/eli/dir/2010/63/oj (accessed September 6, 2024).

[31] J.G. Caporaso, C.L. Lauber, W.A. Walters, D. Berg-Lyons, C.A. Lozupone, P.J. Turnbaugh, N. Fierer, R. Knight, Global patterns of 16S rRNA diversity at a depth of millions of sequences per sample, Proc. Natl. Acad. Sci. U. S. A. 108 Suppl 1 (2011) 4516–4522. 10.1073/pnas.1000080107.

[32] P.D. Schloss, S.L. Westcott, T. Ryabin, J.R. Hall, M. Hartmann, E.B. Hollister, R.A. Lesniewski, B.B. Oakley, D.H. Parks, C.J. Robinson, J.W. Sahl, B. Stres, G.G. Thallinger, D.J. Van Horn, C.F. Weber, Introducing mothur: Open-Source, Platform-Independent, Community-Supported Software for Describing and Comparing Microbial Communities, Appl. Environ. Microbiol. 75 (2009) 7537–7541. 10.1128/AEM.01541-09.

[33] C. Quast, E. Pruesse, P. Yilmaz, J. Gerken, T. Schweer, P. Yarza, J. Peplies, F.O. Glöckner, The SILVA ribosomal RNA gene database project: improved data processing and web-based tools, Nucleic Acids Res. 41 (2013) D590–D596. 10.1093/nar/gks1219.

[34] B.L. Maidak, G.J. Olsen, N. Larsen, R. Overbeek, M.J. McCaughey, C.R. Woese, The RDP (Ribosomal Database Project), Nucleic Acids Res. 25 (1997) 109–110. 10.1093/nar/25.1.109.

[35] R.D. Stedtfeld, X. Guo, T.M. Stedtfeld, H. Sheng, M.R. Williams, K. Hauschild, S. Gunturu, L. Tift, F. Wang, A. Howe, B. Chai, D. Yin, J.R. Cole, J.M. Tiedje, S.A. Hashsham, Primer set 2.0 for highly parallel qPCR array targeting antibiotic resistance genes and mobile genetic elements, FEMS Microbiol. Ecol. 94 (2018) fiy130. 10.1093/femsec/fiy130.

[36] P. Fang, P. Xiao, F. Tan, Y. Mo, H. Chen, U. Klümper, T.U. Berendonk, J. Yang, Biogeographical Patterns of Bacterial Communities and Their Antibiotic Resistomes in the Inland Waters of Southeast China, Microbiol. Spectr. 10 (2022) e0040622. 10.1128/spectrum.00406-22.

[37] Y.-G. Zhu, T.A. Johnson, J.-Q. Su, M. Qiao, G.-X. Guo, R.D. Stedtfeld, S.A. Hashsham, J.M. Tiedje, Diverse and abundant antibiotic resistance genes in Chinese swine farms, Proc. Natl. Acad. Sci. U. S. A. 110 (2013) 3435–3440. 10.1073/pnas.1222743110.

[38] T.D. Schmittgen, K.J. Livak, Analyzing real-time PCR data by the comparative C(T) method, Nat. Protoc. 3 (2008) 1101–1108. 10.1038/nprot.2008.73.

[39] Posit team, RStudio: Integrated Development Environment for R. Posit Software, PBC, Boston, MA., (2024). http://www.posit.co.

[40] H. Wickham, Data Analysis, in: H. Wickham (Ed.), Ggplot2 Elegant Graph. Data Anal., Springer International Publishing, Cham, 2016: pp. 189–201. 10.1007/978-3-319-24277-4_9.

[41] J. Oksanen, G.L. Simpson, F.G. Blanchet, R. Kindt, P. Legendre, P.R. Minchin, R.B. O’Hara, P. Solymos, M.H.H. Stevens, E. Szoecs, H. Wagner, M. Barbour, M. Bedward, B. Bolker, D. Borcard, G. Carvalho, M. Chirico, M. De Caceres, S. Durand, H.B.A. Evangelista, R. FitzJohn, M. Friendly, B. Furneaux, G. Hannigan, M.O. Hill, L. Lahti, D. McGlinn, M.-H. Ouellette, E. Ribeiro Cunha, T. Smith, A. Stier, C.J.F. Ter Braak, J. Weedon, vegan: Community Ecology Package, (2001) 2.6-6.1. 10.32614/CRAN.package.vegan.

[42] S. Atashgahi, R. Aydin, M.R. Dimitrov, D. Sipkema, K. Hamonts, L. Lahti, F. Maphosa, T. Kruse, E. Saccenti, D. Springael, W. Dejonghe, H. Smidt, Impact of a wastewater treatment plant on microbial community composition and function in a hyporheic zone of a eutrophic river, Sci. Rep. 5 (2015) 17284. 10.1038/srep17284.

[43] D. Numberger, L. Ganzert, L. Zoccarato, K. Mühldorfer, S. Sauer, H.-P. Grossart, A.D. Greenwood, Characterization of bacterial communities in wastewater with enhanced taxonomic resolution by full-length 16S rRNA sequencing, Sci. Rep. 9 (2019) 9673. 10.1038/s41598-019-46015-z.

[44] I. Boeraș, A. Burcea, D. Bănăduc, D.-I. Florea, A. Curtean-Bănăduc, Lotic Ecosystem Sediment Microbial Communities’ Resilience to the Impact of Wastewater Effluents in a Polluted European Hotspot—Mureș Basin (Transylvania, Romania), Water 16 (2024) 402. 10.3390/w16030402.

[45] S.A. Wakelin, M.J. Colloff, R.S. Kookana, Effect of Wastewater Treatment Plant Effluent on Microbial Function and Community Structure in the Sediment of a Freshwater Stream with Variable Seasonal Flow, Appl. Environ. Microbiol. 74 (2008) 2659–2668. 10.1128/AEM.02348-07.

[46] B. Drury, E. Rosi-Marshall, J.J. Kelly, Wastewater Treatment Effluent Reduces the Abundance and Diversity of Benthic Bacterial Communities in Urban and Suburban Rivers, Appl. Environ. Microbiol. 79 (2013) 1897. 10.1128/AEM.03527-12.

[47] J. Wang, Y. Li, P. Wang, L. Niu, W. Zhang, C. Wang, Response of bacterial community compositions to different sources of pollutants in sediments of a tributary of Taihu Lake, China, Environ. Sci. Pollut. Res. 23 (2016) 13886–13894. 10.1007/s11356-016-6573-9.

[48] F.-É. Sylvain, A. Holland, S. Bouslama, É. Audet-Gilbert, C. Lavoie, A.L. Val, N. Derome, Fish Skin and Gut Microbiomes Show Contrasting Signatures of Host Species and Habitat, Appl. Environ. Microbiol. 86 (2020) e00789–20. 10.1128/AEM.00789-20.

[49] Y. Luan, M. Li, W. Zhou, Y. Yao, Y. Yang, Z. Zhang, E. Ringø, R. Erik Olsen, J. Liu Clarke, S. Xie, K. Mai, C. Ran, Z. Zhou, The Fish Microbiota: Research Progress and Potential Applications, Engineering 29 (2023) 137–146. 10.1016/j.eng.2022.12.011.

[50] P.S. Kim, N.-R. Shin, J.-B. Lee, M.-S. Kim, T.W. Whon, D.-W. Hyun, J.-H. Yun, M.-J. Jung, J.Y. Kim, J.-W. Bae, Host habitat is the major determinant of the gut microbiome of fish, Microbiome 9 (2021) 166. 10.1186/s40168-021-01113-x.

[51] J. Park, E.B. Kim, Insights into the Gut and Skin Microbiome of Freshwater Fish, Smelt (Hypomesus nipponensis), Curr. Microbiol. 78 (2021) 1798–1806. 10.1007/s00284-021-02440-w.

[52] D. Rosado, P. Canada, S. Marques Silva, N. Ribeiro, P. Diniz, R. Xavier, Disruption of the skin, gill, and gut mucosae microbiome of gilthead seabream fingerlings after bacterial infection and antibiotic treatment, FEMS Microbes 4 (2023) xtad011. 10.1093/femsmc/xtad011.

[53] W. Muziasari, The Resistome of Farmed Fish Feces Contributes to the Enrichment of Antibiotic Resistance Genes in Sediments below Baltic Sea Fish Farms, Front. Microbiol. 7 (2017). 10.3389/fmicb.2016.02137.

[54] M. Ghanbari, W. Kneifel, K.J. Domig, A new view of the fish gut microbiome: Advances from next-generation sequencing, Aquaculture 448 (2015) 464–475. 10.1016/j.aquaculture.2015.06.033.

[55] F.M. Aarestrup, M.E.J. Woolhouse, Using sewage for surveillance of antimicrobial resistance, Science 367 (2020) 630–632. 10.1126/science.aba3432.

[56] P.M.C. Huijbers, C.-F. Flach, D.G.J. Larsson, A conceptual framework for the environmental surveillance of antibiotics and antibiotic resistance, Environ. Int. 130 (2019) 104880. 10.1016/j.envint.2019.05.074.

[57] W. Ahmed, K.A. Hamilton, A. Lobos, B. Hughes, C. Staley, M.J. Sadowsky, V.J. Harwood, Quantitative microbial risk assessment of microbial source tracking markers in recreational water contaminated with fresh untreated and secondary treated sewage, Environ. Int. 117 (2018) 243–249. 10.1016/j.envint.2018.05.012.

[58] L. Opatowski, M. Opatowski, S. Vong, L. Temime, A One-Health Quantitative Model to Assess the Risk of Antibiotic Resistance Acquisition in Asian Populations: Impact of Exposure Through Food, Water, Livestock and Humans, Risk Anal. 41 (2021) 1427–1446. 10.1111/risa.13618.

[59] A.-N. Zhang, J.M. Gaston, C.L. Dai, S. Zhao, M. Poyet, M. Groussin, X. Yin, L.-G. Li, M.C.M. van Loosdrecht, E. Topp, M.R. Gillings, W.P. Hanage, J.M. Tiedje, K. Moniz, E.J. Alm, T. Zhang, An omics-based framework for assessing the health risk of antimicrobial resistance genes, Nat. Commun. 12 (2021) 4765. 10.1038/s41467-021-25096-3.

[60] A.F.C. Leonard, L. Zhang, A.J. Balfour, R. Garside, P.M. Hawkey, A.K. Murray, O.C. Ukoumunne, W.H. Gaze, Exposure to and colonisation by antibiotic-resistant *E. coli* in UK coastal water users: Environmental surveillance, exposure assessment, and epidemiological study (Beach Bum Survey), Environ. Int. 114 (2018) 326–333. 10.1016/j.envint.2017.11.003.

[61] L.E. Furness, A. Campbell, L. Zhang, W.H. Gaze, R.A. McDonald, Wild small mammals as sentinels for the environmental transmission of antimicrobial resistance, Environ. Res. 154 (2017) 28–34. 10.1016/j.envres.2016.12.014.

[62] C. Plaza-Rodríguez, K. Alt, M. Grobbel, J.A. Hammerl, A. Irrgang, I. Szabo, K. Stingl, E. Schuh, L. Wiehle, B. Pfefferkorn, S. Naumann, A. Kaesbohrer, B.-A. Tenhagen, Wildlife as Sentinels of Antimicrobial Resistance in Germany?, Front. Vet. Sci. 7 (2021). 10.3389/fvets.2020.627821.

[63] D. Zeballos-Gross, Z. Rojas-Sereno, M. Salgado-Caxito, P. Poeta, C. Torres, J.A. Benavides, The Role of Gulls as Reservoirs of Antibiotic Resistance in Aquatic Environments: A Scoping Review, Front. Microbiol. 12 (2021). 10.3389/fmicb.2021.703886.

[64] F. Ju, K. Beck, X. Yin, A. Maccagnan, C.S. McArdell, H.P. Singer, D.R. Johnson, T. Zhang, H. Bürgmann, Wastewater treatment plant resistomes are shaped by bacterial composition, genetic exchange, and upregulated expression in the effluent microbiomes, ISME J. 13 (2019) 346–360. 10.1038/s41396-018-0277-8.

[65] D. Leroy-Freitas, E.C. Machado, A.F. Torres-Franco, M.F. Dias, C.D. Leal, J.C. Araújo, Exploring the microbiome, antibiotic resistance genes, mobile genetic element, and potential resistant pathogens in municipal wastewater treatment plants in Brazil, Sci. Total Environ. 842 (2022) 156773. 10.1016/j.scitotenv.2022.156773.

[66] K. Grond, B.K. Sandercock, A. Jumpponen, L.H. Zeglin, The avian gut microbiota: community, physiology and function in wild birds, J. Avian Biol. 49 (2018) e01788. 10.1111/jav.01788.

