## Supplementary Figures for "Fish are poor sentinels for surveillance of riverine AMR"

Germany

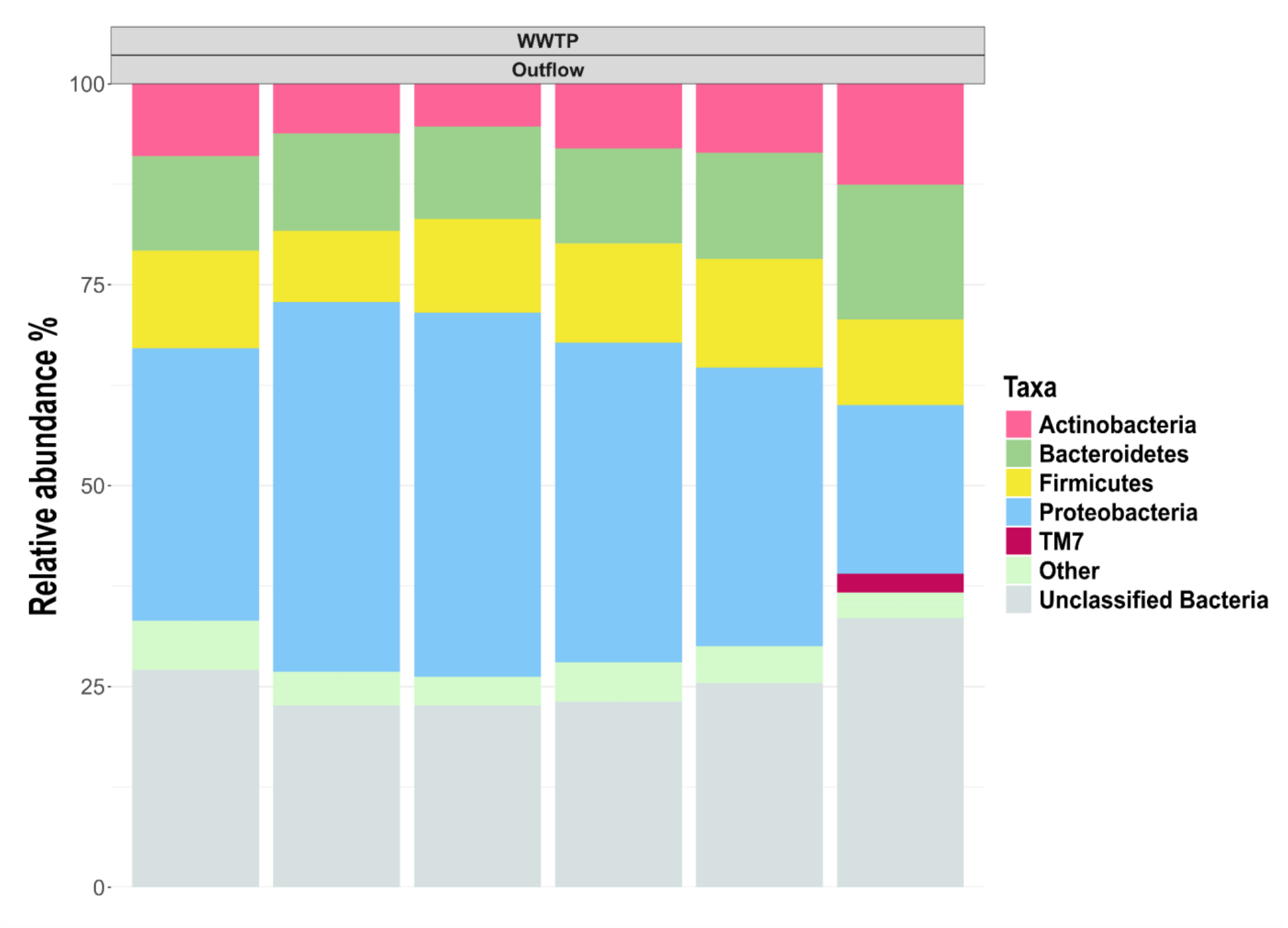


**SI Figure 1: Relative abundance of the phyla dominated in bacterial community composition of the wastewater effluent across six replicate samples. Dominant phyla are defined as those with an average relative abundance of more than 2%. The remaining phyla are grouped as “Other”.**

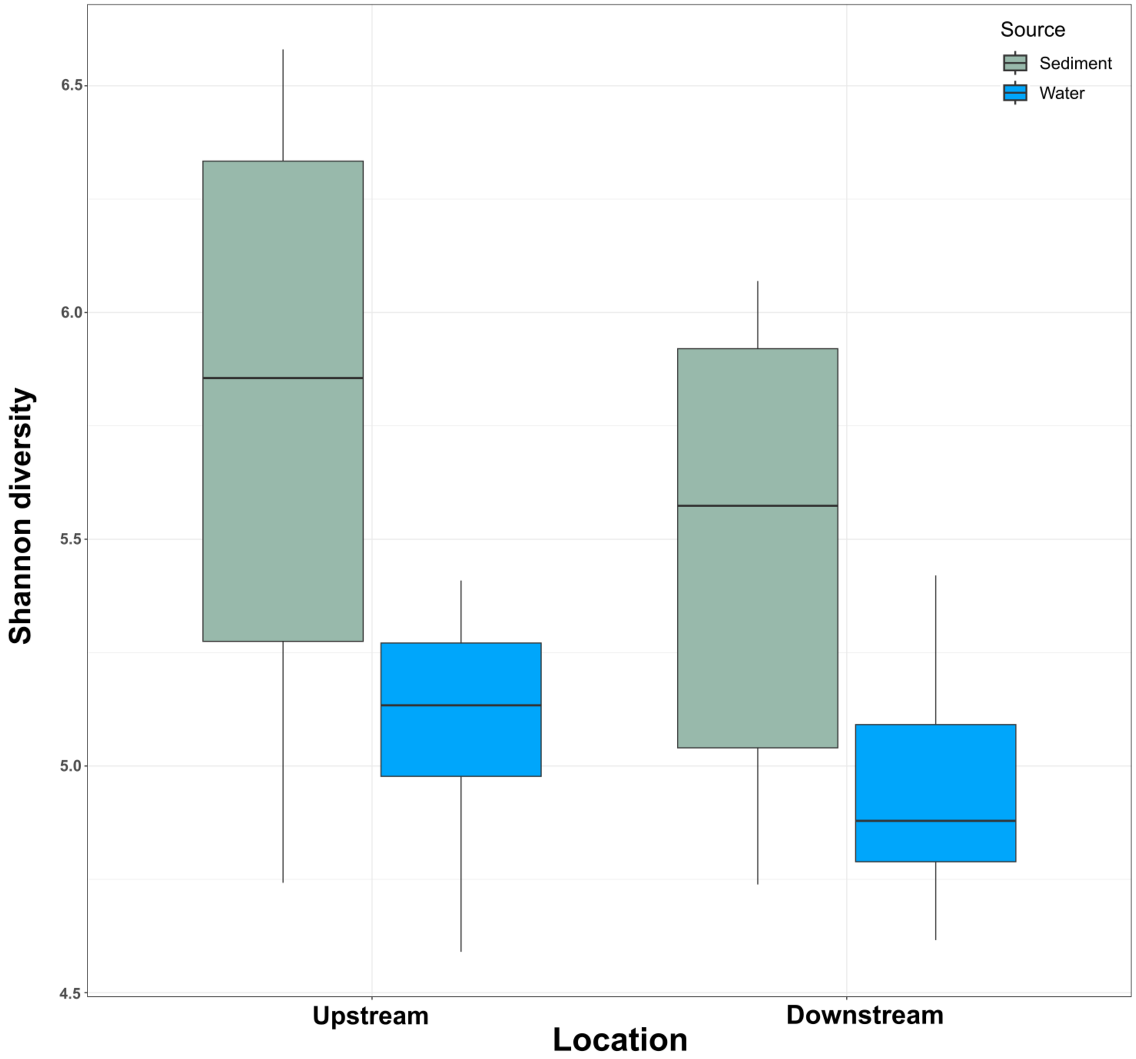


**SI Figure 2. Shannon diversity index for water (A) and sediment (B) samples collected up- and downstream of the WWTP. The boxplots show the distribution of the Shannon index (H) at each location relative to the WWTP. The horizontal line indicates the median diversity value for each location.**

| ARGs | Upstream | Downstream | Outflow | WWTP/Up |
|---|---|---|---|---|
| *blaOXA58* | 0 | 0.00007±0.00003 | 0.00041±0.0001 | ∞ |
| *ermB* | 0.00006±0.00001 | 0.00078±0.0001 | 0.00791±0.0020 | 141.4 |
| *ermF* | 0.000027±0.00002 | 0.00025±0.00005 | 0.00162±0.0004 | 58.3 |
| *tetW* | 0.000067±0.00001 | 0.00033±0.00004 | 0.00246±0.0004 | 36.4 |
| *aac3-VI* | 0.225±0.0425 | 0.2191±0.0313 | 1.81±1.5228 | 8 |
| *qnrS* | 0.00002±0.00001 | 0.000026±0.00001 | 0.00015±0.00002 | 7.3 |
| *blaKPC-3* | 0.000003±0.000004 | 0.000004±0.00001 | 0.00002±0.00002 | 6.4 |
| *dfrA8* | 0.00034±0.00006 | 0.00034±0.00008 | 0.00208±0.0006 | 6.1 |
| *aph6* | 0.00018±0.00004 | 0.00015±0.00002 | 0.00096±0.0002 | 5.3 |
| *vanA* | 0.0326±0.0077 | 0.06076±0.0111 | 0.16516±0.0410 | 5.1 |
| *mphA* | 0.0241±0.0064 | 0.04056±0.0077 | 0.11575±0.0312 | 4.8 |
| *blaVIM* | 0.00016±0.00004 | 0.00012±0.00003 | 0.00078±0.0002 | 4.8 |
| *sul1* | 0.00090±0.0001 | 0.00188±0.0002 | 0.00417±0.0005 | 4.6 |
| *qepA* | 0.09268±0.0194 | 0.1117±0.0191 | 0.35166±0.1806 | 3.8 |
| *aph3-ib* | 0.0819±0.0167 | 0.0651±0.0163 | 0.30316±0.0761 | 3.7 |
| *oqxA* | 0.00045±0.0001 | 0.00016±0.00005 | 0.00165±0.0003 | 3.6 |
| *mcr1* | 0.00541±0.0010 | 0.00368±0.0006 | 0.0178±0.0033 | 3.3 |
| *blaTEM* | 0.00013±0.00006 | 0.00011±0.00005 | 0.00041±0.0001 | 3 |
| *blaKPC-2* | 0.00016±0.00005 | 0.00017±0.00003 | 0.00051±0.0001 | 3 |
| *tetA* | 0.000037±0.00001 | 0.00004±0.00003 | 0.00010±0.00005 | 2.9 |
| *blaCMY-2* | 0.00261±0.0007 | 0.00318±0.0006 | 0.00759±0.0014 | 2.9 |
| *blaCTX-M* | 0.00772±0.0023 | 0.00987±0.0018 | 0.01721±0.0051 | 2.2 |
| *aac(6)-Ib* | 0.00124±0.0003 | 0.0012±0.0002 | 0.00268±0.0008 | 2.2 |
| *blaOXA48* | 0.00697±0.0021 | 0.01280±0.0055 | 0.00941±0.0019 | 1.4 |
| *dfrA1* | 0.000098±0.00004 | 0.00011±0.00004 | 0.00010±0.00002 | 1 |
| *blaNDM* | 0.00397±0.0007 | 0.00299±0.0007 | 0.00191±0.0009 | 0.5 |

**SI Figure 3. Relative abundance of the ARGs in water samples. The numbers represent the average relative abundance of six biological replicates, with standard deviation. The WWTP/Upstream ratio was calculated by dividing the average relative abundance of each gene upstream of the WWTP by the relative abundance of the WWTP effluent. ARGs are presented in descending order of their P >WWTP/Upstream ratio. Ratios, where the ARG was not detected upstream but was detected in the WWTP effluent, are given as ∞. A two-tailed t-test with Bonferroni correction for multiple testing was performed to test the statistical significance of the difference in relative abundance of ARGs up- and downstream of the WWTP. Relative abundance highlighted in green indicates those ARGs with a statistically significant increase of the relative abundance downstream of the WWTP ($P<0.05$). Relative abundance highlighted in purple shows the ARGs with a statistically significant decrease in the relative abundance of the ARGs ($P<0.05$). This supplementary figure provides a detailed representation of the data presented in Fig. 6, particularly regarding the average relative abundance of ARGs per one copy of the 16S rRNA gene, along with standard deviations.**

| ARGs | Trout feces Upstream | Trout feces Downstream | Trout gills Upstream | Trout gills Downstream | Trout skin Upstream | Trout skin Downstream | Water Downstream | Water WWTP/Up |
|---|---|---|---|---|---|---|---|---|
| *blaOXA58* | n.d | n.d | n.d | n.d | n.d | n.d | 0.0000 ± 0.00 | ∞ |
| *ermB* | 0.0008 ± 0.00 | 0.0028 ± 0.00 | 0 ± 0 | 0.0023 ± 0.00 | n.d | n.d | 0.0007 ± 0.00 | 141.39 |
| *ermF* | 0.0018 ± 0.00 | 0.0000 ± 0.00 | n.d | n.d | n.d | n.d | 0.0002 ± 0.00 | 58.29 |
| *tetW* | 0.0000 ± 0.00 | 0.0002 ± 0.00 | n.d | n.d | 0.0020 ± 0.00 | 0 ± 0 | 0.0003 ± 0.00 | 36.39 |
| *aac3.VI* | 0.2361 ± 0.21 | 0.0366 ± 0.02 | 0.5237 ± 0.14 | 0.2615 ± 0.11 | 0.508 ± 0.36 | 0.0415 ± 0.03 | 0.2191 ± 0.03 | 8.04 |
| *qnrS* | 0.0013 ± 0.00 | 0.0001 ± 0.00 | 0.0200 ± 0.01 | 0.0262 ± 0.01 | 0.0230 ± 0.01 | 0.0184 ± 0.01 | 0.0000 ± 0.00 | 7.32 |
| *blaKPC-3* | n.d | n.d | 0 ± 0 | 0.0028 ± 0.00 | n.d | n.d | 0.0000 ± 0.00 | 6.37 |
| *dfrA8* | 0.0043 ± 0.00 | 0.0001 ± 0.00 | 0.0135 ± 0.01 | 0.0069 ± 0.00 | 0.0082 ± 0.01 | 0 ± 0 | 0.0003 ± 0.00 | 6.07 |
| *aph6* | 0.0020 ± 0.00 | 0.0000 ± 0.00 | 0.0188 ± 0.01 | 0.0257 ± 0.02 | 0.0477 ± 0.05 | 0.0027 ± 0.00 | 0.0001 ± 0.00 | 5.28 |
| *vanA* | 0.7029 ± 1.34 | 0.0502 ± 0.03 | 0.8605 ± 0.25 | 1.1115 ± 0.63 | 0.7963 ± 0.42 | 0.0351 ± 0.04 | 0.0607 ± 0.01 | 5.07 |
| *mphA* | 0.1203 ± 0.19 | 0.0179 ± 0.01 | 0.2942 ± 0.08 | 0.3792 ± 0.21 | 0.2363 ± 0.10 | 0.0281 ± 0.03 | 0.0405 ± 0.00 | 4.8 |
| *blaVIM* | 0.0004 ± 0.00 | 0.0000 ± 0.00 | 0.0043 ± 0.00 | 0 ± 0 | n.d | n.d | 0.0001 ± 0.00 | 4.78 |
| *sul1* | 0.0022 ± 0.00 | 0.0007 ± 0.00 | 0.0016 ± 0.00 | 0.0070 ± 0.00 | 0 ± 0 | 0.0098 ± 0.01 | 0.0018 ± 0.00 | 4.59 |
| *qepA* | 0.0752 ± 0.08 | 0.0376 ± 0.03 | 0.2115 ± 0.04 | 0.1887 ± 0.06 | 0.1538 ± 0.07 | 0.0160 ± 0.01 | 0.1117 ± 0.01 | 3.79 |
| *aph3.ib* | 0.7973 ± 1.22 | 0.0278 ± 0.02 | 4.1075 ± 0.87 | 2.39 ± 0.74 | 4.7816 ± 2.97 | 0.425 ± 0.14 | 0.0651 ± 0.01 | 3.7 |
| *oqxA* | 0.0119 ± 0.02 | 0.0001 ± 0.00 | 0.0154 ± 0.01 | 0.0026 ± 0.00 | 0.0169 ± 0.02 | 0 ± 0 | 0.0001 ± 0.00 | 3.62 |
| *mcr1* | 0.0139 ± 0.00 | 0.0017 ± 0.00 | 0.0436 ± 0.01 | 0.0349 ± 0.01 | 0.0694 ± 0.04 | 0.0109 ± 0.01 | 0.0036 ± 0.00 | 3.29 |
| *blaKPC-2* | 0.0072 ± 0.01 | 0.0001 ± 0.00 | 0.0140 ± 0.01 | 0.0038 ± 0.00 | 0.0299 ± 0.04 | 0 ± 0 | 0.0001 ± 0.00 | 3.04 |
| *blaTEM* | 0.0111 ± 0.02 | 0.0020 ± 0.00 | 0.0032 ± 0.00 | 0.0167 ± 0.01 | 0.0198 ± 0.04 | 0.0442 ± 0.08 | 0.0001 ± 0.00 | 3.01 |
| *blaCMY-2* | 0.0429 ± 0.08 | 0.0010 ± 0.00 | 0.0601 ± 0.04 | 0.0719 ± 0.02 | 0.1303 ± 0.10 | 0.0025 ± 0.00 | 0.0031 ± 0.00 | 2.9 |
| *tetA* | 0.0000 ± 0.00 | 0.0000 ± 0.00 | 0.0020 ± 0.00 | 0 ± 0 | n.d | n.d | 0.0000 ± 0.00 | 2.88 |
| *blaCTX-M* | 0.1422 ± 0.30 | 0.0025 ± 0.00 | 0.0687 ± 0.09 | 0.1775 ± 0.07 | 0.1765 ± 0.07 | 0.0064 ± 0.01 | 0.0098 ± 0.00 | 2.23 |
| *aac(6)-Ib* | 0.0087 ± 0.01 | 0.0002 ± 0.00 | 0.0456 ± 0.01 | 0.0259 ± 0.01 | 0.0401 ± 0.01 | 0 ± 0 | 0.0012 ± 0.00 | 2.17 |
| *blaOXA48* | 0.0067 ± 0.00 | 0.0009 ± 0.00 | 0.0144 ± 0.01 | 0.0120 ± 0.00 | 0.0260 ± 0.03 | 0 ± 0 | 0.0128 ± 0.00 | 1.35 |
| *dfrA1* | 0.0002 ± 0.00 | 0.0000 ± 0.00 | 0.0017 ± 0.00 | 0 ± 0 | 0.0020 ± 0.00 | 0.0122 ± 0.02 | 0.0001 ± 0.00 | 1.02 |
| *blaNDM* | 0.0187 ± 0.04 | 0.0003 ± 0.00 | 0.0185 ± 0.01 | 0.0138 ± 0.00 | 0.0528 ± 0.04 | 0 ± 0 | 0.0029 ± 0.00 | 0.48 |

**SI Figure 4: Relative abundance of the ARGs in trout samples. The numbers represent the average relative abundance of the biological replicates, with standard deviation. A two-tailed t-test with Bonferroni correction for multiple testing was performed to test the statistical significance of the difference in relative abundance of ARGs up- and downstream of the WWTP. Relative abundance highlighted in green indicates the ARGs with a statistically significant increase of the relative abundance downstream of the WWTP ($P<0.05$). Relative abundance highlighted in purple is the ARGs with a statistically significant decrease in the relative abundance of the ARGs ($P<0.05$). The column with the relative abundance of the ARGs observed in water downstream of the WWTP indicates the resistance genes, whose increase in relative abundance was impacted by the effluent. Ratios, where the ARG was not detected in water samples upstream of the WWTP but was detected in the WWTP effluent, are given as ∞. This supplementary figure provides a detailed representation of the data presented in Fig. 7, particularly regarding the average relative abundance of ARGs per one copy of the 16S rRNA gene, along with standard deviations.**

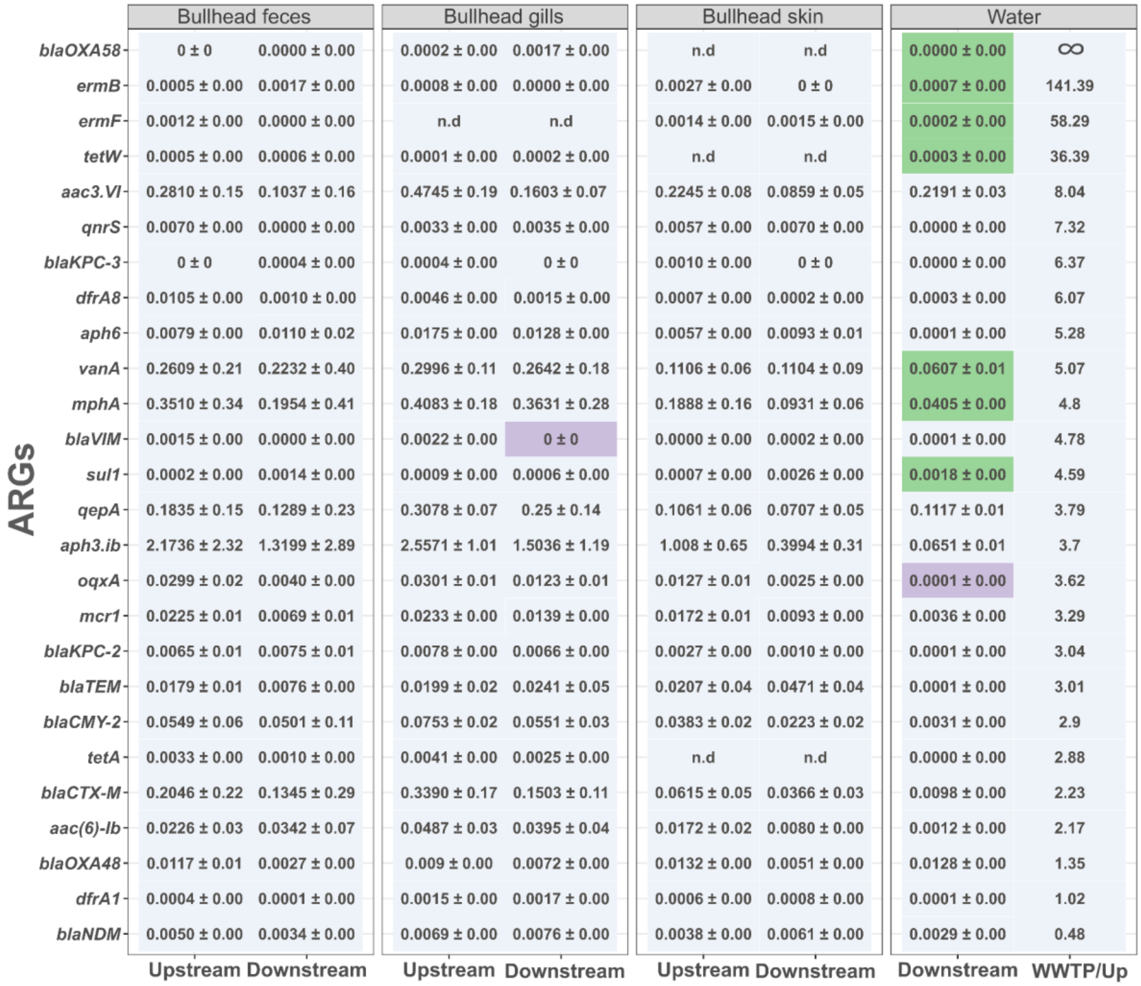


**SI Figure 5. Relative abundance of the ARGs in bullhead samples. The numbers represent the average relative abundance of the biological replicates, with standard deviation. A two-tailed t-test with Bonferroni correction for multiple testing was performed to test the statistical significance of the difference in relative abundance of ARGs up- and downstream of the WWTP. Relative abundance highlighted in green indicates the ARGs with a statistically significant increase of the relative abundance downstream of the WWTP (P<0.05). Relative abundance highlighted in purple is the ARGs with a statistically significant decrease in the relative abundance of the ARGs (P<0.05). The column with the relative abundance of the ARGs observed in water downstream of the WWTP indicates the resistance genes, whose increase in relative abundance was impacted by the effluent. Ratios, where the ARG was not detected in water samples upstream of the WWTP but was detected in the WWTP effluent, are given as ∞.**
